# Variable visual errors suppress implicit motor adaptation among older adults

**DOI:** 10.64898/2026.11.06.764937

**Authors:** Yan Zhou, Iulia Andrews, Alicia Sang, Joaquin Torres, Olivia A. Kim

**Author notes:** **Corresponding Author:** Olivia A. Kim. Department of Industrial/Organizational Psychology, New York University, New York, NY 10003, USA. Adams Clinical, Boston, MA 02116, USA. **Contributions: *Yan Zhou:*** Formal Analysis, Investigation, Visualization, Writing – original draft; ***Iulia Andrews:*** Investigation, Writing – review & editing; ***Alicia Sang:*** Investigation, Writing – review & editing; ***Joaquin Torres:*** Investigation, Writing – review & editing; ***Olivia Kim:*** Conceptualization, Formal Analysis, Funding Acquisition, Investigation, Methodology, Project administration, Resources, Software, Supervision, Visualization, Writing – original draft, Writing – review & editing. **Funding Sources:** This research did not receive any specific grant from funding agencies in the public, commercial, or not-for-profit sectors.

## Abstract

Age-related changes in motor control networks manifest in a highly task-dependent manner. Although older adults can display enhanced implicit sensorimotor adaptation, this effect is inconsistent across perturbation types (*e.g.*, visuomotor vs. force-field perturbations). Because these perturbations induce different errors to drive learning, their distinct effects on learning suggest that specific properties of the error signal modulate the extent of adaptation among older adults. Here, we investigated whether age-related hyper-adaptation is modulated by error variability. More specifically, we manipulated the consistency of error direction during an error-clamp visuomotor adaptation task in both younger and older adults. While consistent errors elicited enhanced implicit adaptation in older adults relative to younger controls, variable errors did not. These findings reveal that error consistency is a critical prerequisite for age-related enhancements in implicit motor learning. Rather than exhibiting unconstrained hyper-adaptation, older adults maintain an adaptive error-sensitivity mechanism that prevents hyper-recruitment of implicit systems in noisy motor environments.

**Graphical Abstract:** 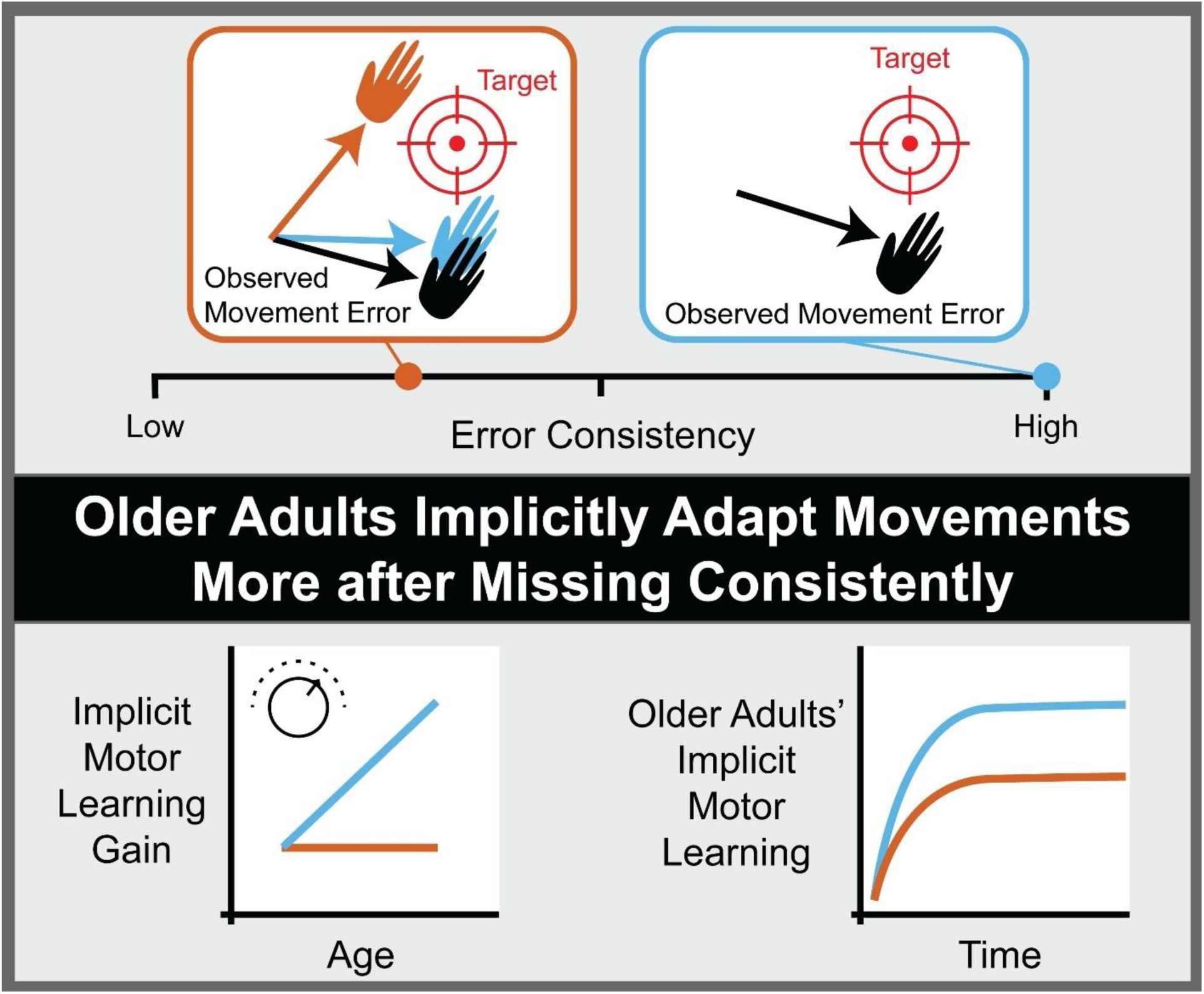

## 1. Introduction

Brain regions vital for motor learning and control, such as the cerebellum and motor cortices, shrink by up to 40% across the lifespan, likely contributing to age-related differences in our movements (Andersen et al., 2003; Arleo et al., 2024; C. Clark and L. Taylor, 2011; d’Oleire Uquillas et al., 2026; Mooney et al., 2022). However, age-related motor changes reported in the literature are diverse (Berghuis et al., 2017; Clayton et al., 2024; Heuer and Hegele, 2014; Rasman et al., 2023; Rueda-Delgado et al., 2019; Ruitenberg et al., 2023; Seidler, 2006; Vachon et al., 2020; Wolpe et al., 2020; Zvornik et al., 2024). This complexity creates a need for studies that utilize a task with known neural and cognitive substrates, as such work will contribute to a knowledge base that supports better characterization and understanding of the richness of the aging motor system (Zapparoli et al., 2022).

Presenting an excellent opportunity to parse age-related changes, implicit sensorimotor adaptation tasks are cerebellar-dependent (Krakauer et al., 2019; Tzvi et al., 2022; Manto et al., 2023; Hagen and Fling, 2026; Morehead et al., 2017). Such implicit adaptation proceeds below the level of conscious awareness and contributes to overall learning during sensorimotor adaptation (Mazzoni and Krakauer, 2006; Taylor et al., 2014; McDougle et al., 2015; Krakauer et al., 2019; Hart et al., 2024).

Recent reports have put forth contradictory narratives about age-related changes in implicit adaptation. Studies utilizing error-clamp (EC) visuomotor adaptation tasks that isolated implicit learning highlight that adults over age 58-65 can show up to twice as much adaptation as adults between the ages of 18-35 (Cisneros et al., 2026; Vandevoorde and Orban de Xivry, 2019). This suggests that the implicit process gains the capacity to learn more as we age. However, despite the consensus that learning in force-field tasks typifies implicit adaptation, older adults show neither an enhancement nor a deficit in adaptation of reaches in a force-field (Kim et al., 2021; Hermans et al., 2025). Taking these findings into account, it is clear that implicit adaptation as a whole is not simply amplified among older adults. Age-related modulation of adaptation must depend on something that differs between these distinct sensorimotor tasks.

One key difference between previously deployed EC paradigms and force-field tasks lies in the consistency of the error that drives sensorimotor learning. Adaptation in both tasks is caused by mismatches between the expected and observed consequences of movement (sensory prediction errors, SPEs; Therrien and Wong, 2022). These SPEs include visual errors in which a cursor representing the participant’s hand follows a different trajectory than the participant intended. This is the predominant error in EC tasks, for which the experimenter specifies the direction of visual feedback independent of participants’ actual movements (Morehead et al., 2017). This allowed experimenters to present uniform visual errors during studies that evoked an age-difference in EC-evoked adaptation (**Fig. 1**, top; Vandevoorde and Orban de Xivry, 2019; Cisneros et al., 2026). During force-field driven learning, however, visual error can be more variable. Variability arises in these tasks because they rely on a robot pushing or pulling the moving limb to induce SPEs, and participants learn to apply their own forces to counteract the robot (Kim et al., 2021). This learning involves some deliberate trial-and-error that can result in overcompensation for the force-field, rendering the exact mismatch between the actual and intended visual consequences of movement variable during early adaptation (**Fig. 1**, bottom; Schween et al., 2020). It may be that this error variability limited age-related hyper-adaptation during force-field tasks, whereas hyper-adaptation was facilitated by the highly consistent visual error in EC tasks.

**Figure 1.**
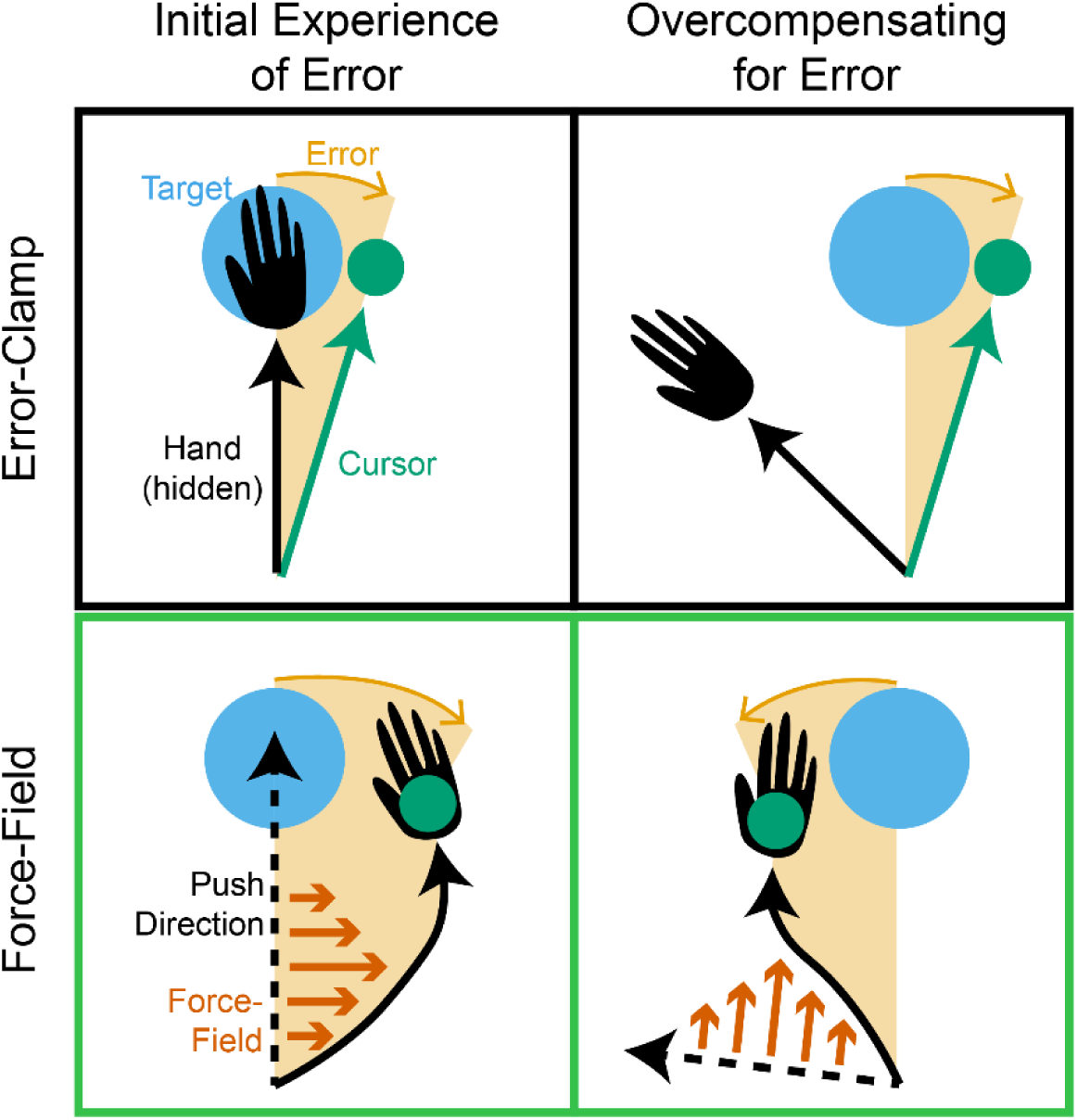
Possible error directions during error-clamp and force-field adaptation tasks. Schematics of hand movement vectors (grey), visual feedback (green) vectors, and visual error (orange) during error-clamp (top) and force-field adaptation (bottom) trials. (Left) Early experiences of errors caused by clockwise perturbations of movements made directly towards the target. The force-field applied is shown in dark orange, and the direction in which participants plan to push is shown with a black dashed line. (Right) Movements made and errors experienced on a trial where a participant overcompensates for the perturbation. For the error-clamp trials, the visual feedback is identical in both conditions. For the force-field trials, the visual feedback is contingent upon the way the participant pushes during their movement. In the example shown here, cursor appears on the opposite side of the target as the initial condition, meaning that the error is in the opposite direction from the perturbation.

Consistency in error direction is critical during sensorimotor adaptation, and there is extensive neural machinery devoted to its processing. For example, error direction is differentially encoded in the cerebellum and dictates the way that learning proceeds (Medina, 2011; Herzfeld et al., 2018; Spampinato and Celnik, 2021; Streng et al., 2022; Shadmehr and Fakharian, 2026). As a consequence, learning is affected by participants’ recent histories of experience with variable error (Albert et al., 2021; Avraham and Ivry, 2025). Given that the history of error direction influences the extent of learning and that reports on age-differences in the amount of learning achieved vary between EC and force-field tasks, variability in experimental control of error direction may contribute to discrepancies in reports on age-differences in sensorimotor adaptation.

Testing whether age-related enhancement of cerebellar-dependent implicit learning depends on exposure to a consistent visual error, we systematically manipulated errors presented to older (65+ years) and younger adults (18-35 years) in a series of experiments employing EC visuomotor adaptation tasks. While EC tasks cannot mimic all kinds of error presented during force-field paradigms, we applied ECs because they provide complete control over visual error. As a result, our experiments directly tested whether variability in visual error alone is sufficient to influence implicit adaptation among older adults.

After replicating findings of enhanced adaptation among older adults in response to uniform errors (Experiment 1), we observed that adaptation did not vary between age groups when error directions were pseudorandomized and fully counterbalanced across training (Experiment 2). We also demonstrated that age-differences in adaptation to a uniform error were blunted following exposure to salient errors in the opposite direction (Experiment 3). Based on these results, we argue that variability in visual error is a crucial mediator of age-related changes in visuomotor adaptation, suggesting that older adults’ implicit systems may selectively adapt more in response to consistent visual errors without overcompensating for random environmental noise.

## 2. Methods

### 2.1. Participants and Selection Criteria

We recruited younger adults (aged 18–35) and older adults (aged 65+) from the Bates College and South-Central Maine communities for in-person testing (Experiments 1 and 2a), and throughout the United States via Prolific (London, UK) for online testing (Experiments 2b and 3). In-person participants received an honorarium of $20/hour; online participants were compensated between $12 and $20/hour per platform guidelines. All procedures were approved by the Bates College Institutional Review Board. All participants provided informed consent.

Sample size targets were determined via a priori power analysis in G*Power (v3.1.9.7; Kiel, Germany). Based on effect sizes from related visuomotor paradigms (e.g., Exp. 4 in (Vandevoorde and Orban de Xivry, 2019), a minimum of 30 participants per group was required to achieve a power of 0.80. To accommodate the inherently noisier data collection typical of online crowdsourcing, the target was increased to 50 completed datasets per group for Experiments 2b and 3. Details about sample sizes, attrition, and final demographic characteristics across all experiments are consolidated in **Table 1**. Handedness across the online cohorts (Experiments 2b and 3) was predominantly right-handed. Participant counts matching dominant hand to the hand used during the task are provided in **Supplementary Table S1**.

**Table 1.**
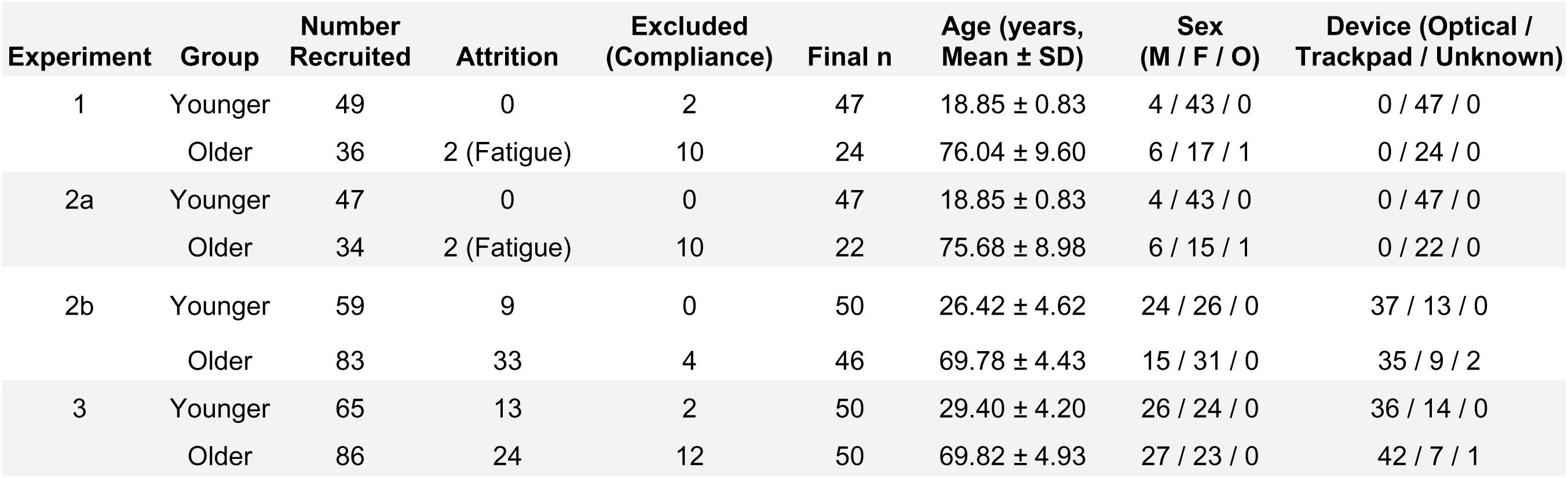
Participant quantities and demographic details.

### 2.2. Exclusion Criteria

To isolate implicit adaptation, which features characteristically slow, gradual adjustments in reach trajectory, from explicit cognitive re-aiming strategies marked by abrupt shifts (Bond and Taylor, 2015; McDougle et al., 2015), we instructed participants to do their best to ignore the cursor direction and aim directly towards the target. However, some participants did not follow these instructions, instead making large, abrupt changes in their movements. To systematically detect and filter out these participants from our dataset, we implemented an automated trial-by-trial filter. Participants were operationalized as having deployed an explicit re-aiming strategy and were excluded from final analysis if they shifted their reach direction by more than 15° on any single trial and sustained that shift such that their mean movement angle across the subsequent 10 trials differed significantly from the preceding 10 trials. We selected 15° as a threshold because this number is substantially greater than the standard amount of variability observed during center-out reaching tasks and likely represents a deliberate adjustment in movement direction (Pekny et al., 2015). This programmatic threshold isolated genuine strategic shifts from isolated exploration/mistake trials. The performance of exemplar participants who were included or excluded from the final are shown in **Supplementary Figure 1**.

### 2.3. Visuomotor Adaptation Paradigm

#### 2.3.1. Apparatus and Core Trial Structure

All experiments utilized a custom browser-based center-out reaching task programmed in JavaScript and hosted on a JATOS server at Bates College. In-person participants (Experiments 1 and 2a) completed the task using a standardized Dell Latitude 5440 laptop and trackpad. Online cohorts (Experiments 2b and 3) utilized their native hardware environments. Based on our and others’ previous work, we do not expect the hardware used to have affected our results (Kim et al., 2022; Tsay et al., 2021).

Each trial progressed dynamically through a sequence. First, the participant guided a white, 2-pixel-radius cursor into an empty central starting circle on a black background. Upon entry, the circle filled grey to confirm positioning, and a randomized pre-stimulus delay (500-1000 ms) began. If the participant moved the cursor out of the center before the target appeared, this phase re-started. After the pre-stimulus delay, a single blue, 8-pixel diameter target appeared 200 pixels away. To eliminate target-location learning biases, the target was locked to one of four off-cardinal locations, randomized across participants. Participants sliced rapidly through the target after it appeared. The moment the cursor exceeded 200-pixels from the center (target distance), visual feedback stopped updating. If visible on the trial, the cursor remained displayed at the location it crossed the 200-pixel boundary for 500 ms. Then, the cursor was automatically reset at a random position within 20 pixels of the center, requiring an active return movement to reset the workspace for the next trial.

To preserve movement fidelity, online kinematic rules were strictly enforced via automated warnings and penalty timeouts. If a movement took longer than 200 ms to span the center-to-target distance, a speed warning and a 2-second timeout penalty were applied to suppress online error correction. To encourage participants to try to move their mouse towards the displayed target, movements deviating more than 60° from the target axis triggered a directional warning and a 2-second timeout. This 60° threshold exceeds the typical range of implicit adaptation and should not have punished participants following instructions in our task (Al-Fawakhiri et al., 2023; Morehead et al., 2017). Hesitation longer than 15 seconds to initiate a trial triggered a visual prompt without a timeout penalty.

#### 2.3.2. Visual Feedback Conditions

We manipulated cursor behavior through three distinct kinds of visual feedback. On trials with veridical feedback (veridical FB), the cursor was displayed at the location of the mouse’s true digital coordinates. On trials with no feedback (no-FB), the cursor was entirely masked after the participant moved the cursor outside of the center starting location towards the target, providing no positional or terminal error cues. Finally, on trials with error-clamped feedback (EC FB), the cursor moved at an invariant angular trajectory relative to the target, completely decoupled from the participant’s physical hand movement. The hand distance from center dictated cursor distance, but the angle was fixed. In the text, “+45° EC” refers to a trial where the cursor was shown moving in a direction 45° counterclockwise of the target, and a “-45° EC” refers to a trial where the cursor was shown moving 45° clockwise of the target. The phrase “45° EC” refers to blocks with a 45° EC in a consistent direction for a given participant.

### 2.4. Task Structure

#### 2.4.1. General Tutorial

The experiment began with a brief, interactive tutorial phase with veridical feedback. During this tutorial, participants learned that the computer mouse controlled the movement of the cursor, that they needed to move to the central starting location to make a target appear, and that they needed to slice through the target to progress through the experiment. The experiment did not begin in earnest until the participant successfully completed 3 trials without receiving a speed warning or a directional warning (see Section 2.3.1 above for more details on trial structure and warnings).

#### 2.4.2. Error-Clamp Tutorial

To guarantee instruction comprehension before entering the critical EC blocks, participants completed a 2-trial tutorial at the end of the baseline phase. During this sequence, they were explicitly told before each trial that the cursor would not follow their movements, and that they should move directly to the target and simply observe what the cursor does. To make the presence of a perturbation obvious, these trials involved 90° ECs. To limit the impact of these tutorial trials on performance in the rest of the task, the target was presented straight ahead of the starting location, which lies far enough from the training target locations to be outside of the typical range of generalization for implicit adaptation (Schween et al., 2018). Additionally, the EC direction was counterbalanced across trials to minimize adaptation during the EC tutorial. Participants then completed a two-choice forced alternative question detailing the instructions (“Press Z if you should do your best to make the cursor hit the target”, “Press X if you should do your best to ignore the cursor and move your real mouse to the target”). The task remained locked until the correct answer (X) was selected. Then, participants received one more text reminder of the task instructions.

#### 2.4.3. Experimental Schedules

The sequence of blocks varied for each experiment to measure implicit adaptation in response to different error schedules.

##### 2.4.3.1. Experiment 1

Experiment 1 consisted of 250 total trials. A baseline block, following standard protocols, consisted of 24 veridical FB trials followed by 24 no-FB trials (Section 2.3.2). Then, 2 EC tutorial trials (Section 2.4.2) preceded a 100-trial learning block consisting of 45° EC trials, with the EC direction randomized between participants. The experiment ended with 100 washout trials, all with 0° ECs applied. Due to fatigue-related attrition among some older adults during the washout block, data analysis was conducted across epochs where data across all participants remain fully populated (up to the 57th trial of washout).

##### 2.4.3.2. Experiment 2

Experiment 2a immediately followed Experiment 1. It consisted of 120 trials with +/-5.25° ECs. Trials were counterbalanced such that each EC direction (+: counterclockwise, -: clockwise) appeared 5 times in every 10 trials. The target appeared on the opposite side of the workspace as in Experiment 1.

Experiment 2b was similar to Experiment 2a, except that the EC trials immediately followed a baseline block (25 veridical feedback trials and 25 no feedback trials) and 4 EC tutorial trials.

During half of the trials in Experiments 2a and 2b, the target was re-centered from its original location to the cursor location displayed during the EC FB (“target jump” manipulation as in (Al-Fawakhiri et al., 2023; Tsay et al., 2022)). Because these target jump trials were intended to address a distinct research question and because of ambiguity in whether older adults followed EC instructions to aim to the original target location despite the jump, target jump trials were excluded from the primary analyses presented here (**Fig. 3B**, **3D**). Exploratory analyses including data from target jump trials did not qualitatively affect the results of our statistical analyses or our conclusions.

##### 2.4.3.3. Experiment 3

Experiment 3 began with a baseline block (10 veridical feedback, 10 No-FB) followed by 2 EC tutorial trials. Then, we presented a learning block with 60, 45° EC trials followed by 10 No-FB trials, with No-FB trials included as a window for the measurement of implicit adaptation without additional error presentation. This was followed by a washout block involving 30 trials with veridical feedback. A second learning block was then presented, including the same 60 trials with a 45° EC and 10 trials No-FB trials.

Before the washout block, participants were instructed that the cursor would once again follow their movements and they should try to get the cursor on the target. Before the second learning block participants were informed that the cursor would try to trick them again and they should do their best to ignore the cursor while getting the computer mouse on the target.

### 2.5. Data Analysis

#### 2.5.1. Quantification of Movement Kinematics

Raw data were processed using Python 3.0 (packages: pandas, numpy). The primary dependent variables were calculated from the coordinates of each movement.

Movement direction (reach angle) was calculated as the angular vector of the mouse coordinate at the moment it first reached or exceeded 200 pixels from the center starting location (target distance). Angles are reported relative to the target, with a movement directly on target having a movement angle of 0° and with movement angles counterclockwise of the target having a positive sign.

Reaction time (RT) was calculated as the latency between the target appearance and the cursor exiting the center starting location.

Movement duration was defined as the elapsed time between the cursor exiting the center starting location and exceeding 200 pixels from the center (target distance).

#### 2.5.2. Statistical Analysis

Inferential statistics were conducted in R. Structural changes across time blocks and between-group demographic variances (Young vs. Older) for Experiments 1 and 3 were analyzed using two-way mixed repeated-measures ANOVA via the afex::aov_ez library and function.

Violations of sphericity were automatically adjusted utilizing Huynh-Feldt corrections, with adjusted p-values noted throughout the text. Effect size bounds for main omnibus tests are expressed as partial eta-squared and converted to *Cohen’s F* via the effectsize package. Pairwise post-hoc decompositions were conducted via emmeans, applying Benjamini-Hochberg False Discovery Rate (FDR) adjustments to correct for multiple comparisons. Post-hoc effect magnitudes are reported as *Cohen’s d*.

## 3. Results

To evaluate the contribution of error history to differences in implicit adaptation between older adults (65+ years old) and younger adults (18-35 years old), we first confirmed that older adults show greater adaptation after a block of unidirectional error-clamps (ECs). Then, in Experiment 2, we fully counterbalanced bidirectional error-clamps throughout training and observed no age-differences in single-trial adaptation (STA). STA evokes small behavioral changes in response to single errors, leaving two possible explanations for the lack of age-differences in Experiment 2: 1) an effect of error history or 2) a floor effect. To discriminate between these possibilities, Experiment 3 tested whether a history of blocks of salient bidirectional errors affected age-differences in accumulated adaptation. Together, the results from these three experiments show implicit adaptation patterns among older adults that resemble those of younger adults when there is a recent history of variable errors.

### 3.1. Unidirectional errors cause age-differences in adaptation

Experiment 1 sought to replicate and qualify the finding that implicit adaptation to ECs is enhanced among older adults. To this end, older and younger adults (*n_older_* = 24, *n_younger_* = 47) completed a computer-based visuomotor adaptation task. The task employed standard protocols for establishing baseline movement features, and then imposed unidirectional 45° errors via a block of EC trials (**Fig. 2A**), similar to previous studies (Cisneros et al., 2026; Vandevoorde and Orban de Xivry, 2019). To extend this research by beginning to characterize the longevity of age-differences in adaptation, we then imposed a 0° EC washout block, during which participants observed no visual errors. The absence of visual errors isolated forgetting as the sole driver of further adaptation (Tanaka et al., 2009), allowing for the washout of short-term adaptation memories and the measurement of longer-lasting adaptation memories.

**Figure 2.**
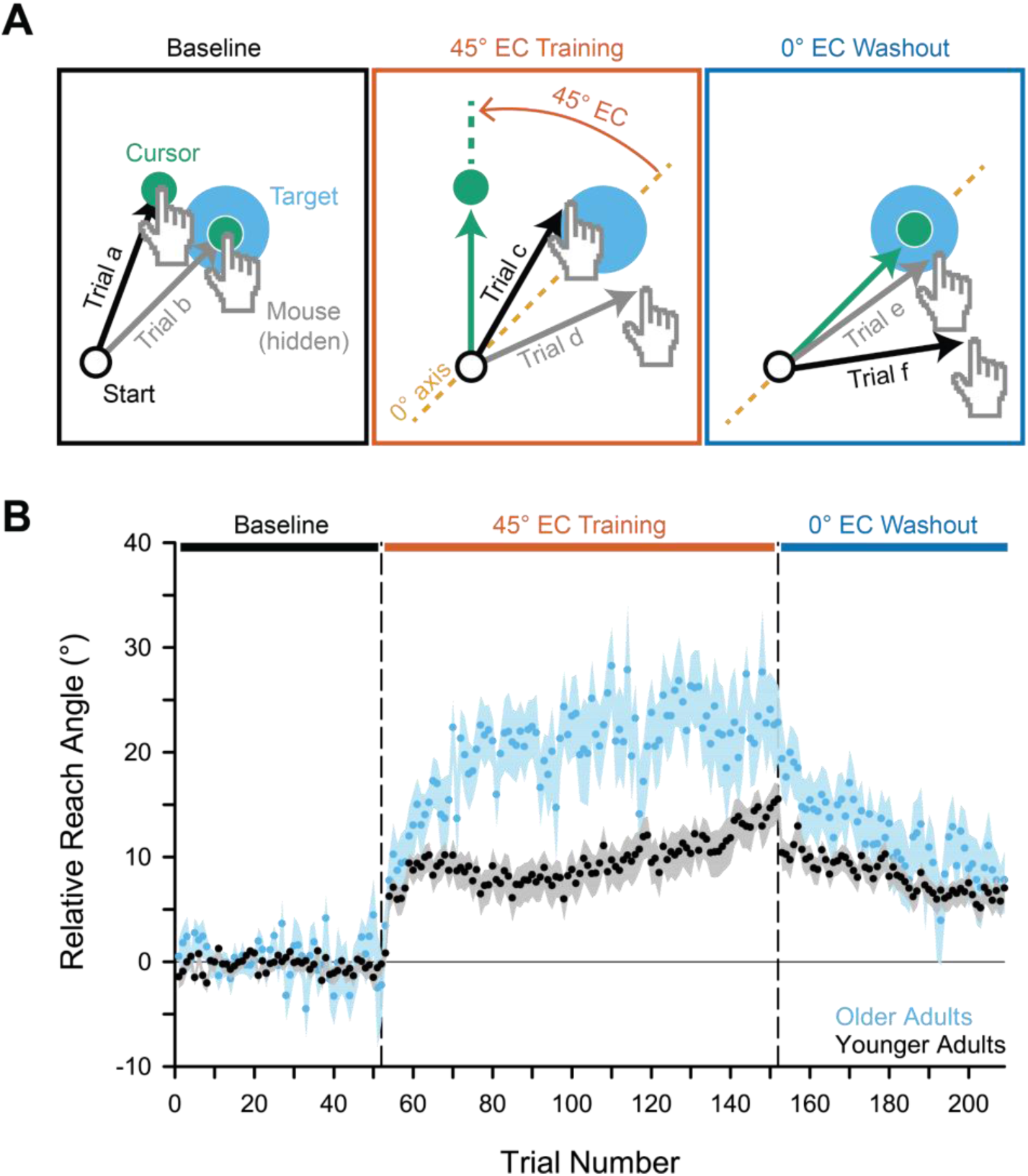
Implicit adaptation during blocked EC training with unidirectional errors. **A.** Schematics indicating the feedback provided during the different phases of the experiment. The borders around each schematic are colored to indicate phase corresponding to the bars in panel **B**. The cursor is shown in green and the workspace background is shown in white for illustrative purposes, but they were white and black, respectively, during the task. **B.** Scatter plot showing the mean ± SEM movement direction across participants. Movement direction is shown relative to the target and relative to the direction of the EC during the main training phase, with positive relative movement direction values corresponding to the direction of movement opposite the direction of the error (*i.e.,* positive values indicate clockwise movements when the EC was in the counterclockwise direction). Data from older adults are shown in blue, and data from younger adults are shown in black. *Abbreviations*: EC: error-clamp, FB: visual feedback, SEM: standard error of the mean; wash: washout.

Participants adapted their movement angles gradually (main effect of Timepoint: *F(3, 207)* = 64.94, *p_corrected_* = 3.62 x 10^-22^, *η^2^_p_* = 0.48, *Cohen’s F* = 0.97) and in a manner that differed between age groups (main effect of Age Group: *F(1, 69)* = 6.17, *p* = 0.01, *η^2^_p_* = 0.08, *Cohen’s F* = 0.30; **Fig. 2B**). These age-differences emerged over the course of the task (Age x Timepoint interaction: *F(3, 207)* = 5.13, *p_corrected_* = 0.006, *η^2^_p_*= 0.07, *Cohen’s F* = 0.27). During the baseline block, both age groups’ movements’ deviations from the target direction did not differ from 0° (mean ± standard error of the mean: 0.22° ± 0.23°; *t(69)* = 1.37, *p_adjusted_*= 0.17). By the end of the 45° EC block, older adults’ movement angles adapted away from the target opposite the direction of the EC (23.59° ± 3.60°). Adaptation among younger adults (13.75° ± 1.80°) was significantly smaller (*t(69)* = 2.74, *p_adjusted_* = 0.008, *Cohen’s d* = 0.66). This group difference was sustained early in the 0° EC washout block (*t(69)* = 2.59, *p_adjusted_* = 0.006, *Cohen’s d* = 0.69), replicating previous findings that a 45° EC block evokes more adaptation in older adults relative to younger adults. Age-differences in adaptation subsided after extended exposure to the 0° EC (*t(69)* = 0.56, *p_adjusted_* = 0.57), with movement angles in both groups appearing to stabilize around 7.01° ± 1.25° relative to the target.

In summary, adaptation is enhanced among older adults during exposure to the unidirectional errors in a 45° EC block, but this enhancement subsides after a period without visual errors in a 0° EC block. This suggests that age-related enhancement of adaptation may depend on continuous exposure to a consistent, nonzero EC.

### 3.2. Fully counterbalanced bidirectional errors do not cause age-differences in adaptation

To test the extent to which age-related enhancement of adaptation depends on exposure to a consistent, nonzero EC, we ran two follow-up experiments. In our first follow-up experiment (Experiment 2a), we continued working with participants (*n_older_* = 22, *n_younger_* = 47) from Experiment 1 to measure adaptation to ECs with randomly varying directions (**Fig. 3A**). Because this pattern of errors causes adaptation that sums destructively and results in 0° of net adaptation (Sheahan et al., 2016), Experiment 2a quantifies adaptation in terms of STA, which we have previously used to measure implicit adaptation in response to single ECs (Al-Fawakhiri et al., 2023; Kim et al., 2022). To maximize the signal-to-noise ratio for detection of any potential age-differences in STA, which tends to be small in amplitude, we presented 5.25° ECs that we previously noted evoked maximal STA (Al-Fawakhiri et al., 2023).

**Figure 3.**
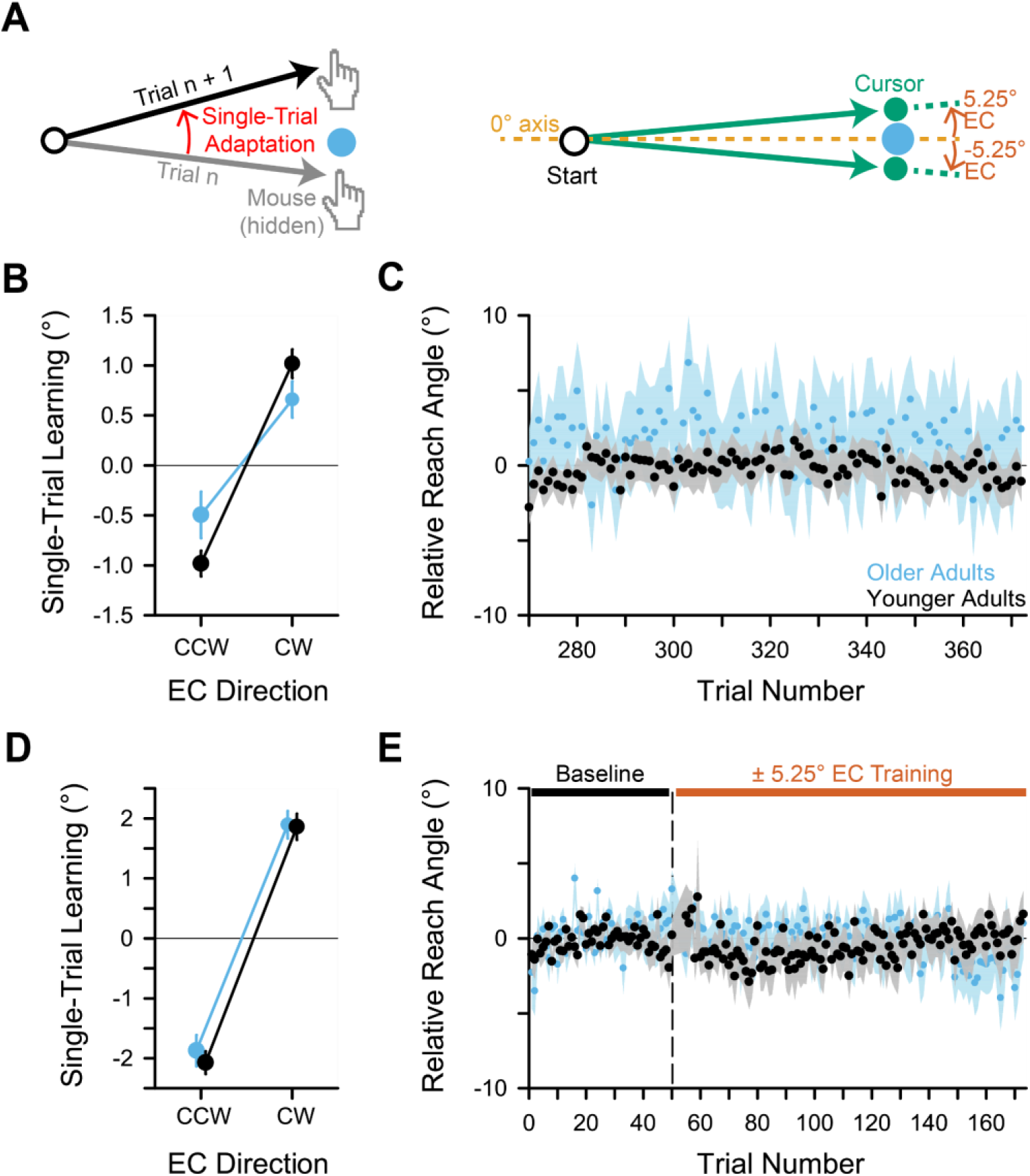
Implicit adaptation to ECs of randomly varying direction. **A.** Schematic illustration of STA calculation and the EC feedback during Experient 2. **B.** Scatter plot showing the mean ± SEM adaptation in movement direction in response to single ECs across participants (STA). Positive values correspond to clockwise changes in movement direction and negative values correspond to counterclockwise changes. Data from older adults are shown in blue, and data from younger adults are shown in black. **C**. Scatter plot showing the mean ± SEM movement direction across participants. Movement direction is shown relative to the target and relative to the direction of the 45° EC during Experiment 1, with positive relative movement direction values corresponding to the direction of movement opposite the direction of the error (*i.e.,* positive values indicate clockwise movements when the EC was in the counterclockwise direction). **D-E.** As in **B-C**, but pertaining to Experiment 2b. *Abbreviations*: B: baseline, EC: error-clamp, SEM: standard error of the mean, STA: single-trial adaptation.

To access a clean slate for new learning, we displayed the target for Experiment 2a on the opposite side of the workspace from the target shown in Experiment 1. Moving the target in this way shifts participant movement angles outside the range of both the plan-based and biomechanical reference frames in which implicit adaptation generalizes (Schween et al., 2018; Poh et al., 2022). Therefore, the behavior measured in Experiment 2a should not have been directly influenced by adaptation that occurred in Experiment 1. Across the experiment, participants exhibited significant STA, with 0.86° ± 0.11° of adaptation opposite the direction of the preceding EC (main effect of EC Direction: *F(1, 67)* = 50.26, *p* = 1.05 x 10^-9^, *η^2^_p_* = 0.43, *Cohen’s F* = 0.87; *t(67)* = -7.10, *p* = 1.05 x 10^-9^, *Cohen’s d* = -1.73; **Fig. 3B**). The magnitude of STA was similar between age groups (main effect of Age Group: *F(1, 67)* = 0.37, *p* = 0.54), although there was a trend towards an interaction between the effects of Age Group and EC Direction (*F(1, 67)* = 3.61, *p* = 0.06). These results suggest that the task paradigm applied for Experiment 2a may not evoke adaptation that differs between younger and older adults, even though the same set of younger and older adults exhibited differences in adaptation in response to a different pattern of ECs in Experiment 1 (**Fig. 2**). However, the trend towards an interaction effect of Age Group and EC Direction makes these results difficult to interpret. Difficulties in interpretation also arise upon examination of movement angles.

Although movement angles were stable and directed towards the target throughout Experiment 2a (mean ± SEM movement angle: -0.13° ± 1.30°, main effect of Timepoint: *F(1, 67)* = 0.63, *p* = 0.43), they tended to be numerically higher among older adults (**Fig. 3C**). This effect was not statistically significant (main effect of Age Group: *F(1, 67)* = 1.83, *p* = 0.18; Age Group x Timepoint interaction: *F(1, 67)* = 0.66, *p* = 0.42), but this could be because our study was not sufficiently powered to detect Experiment 1’s small, carryover effects on movement angle in Experiment 2a.

Because an elevated movement angle during the Experiment 2a would be consistent with residual adaptation from Experiment 1 and because such carryover effects may have undermined our ability to detect any age-differences in STA by changing the baseline relative to which STA was measured, we conducted Experiment 2b. In this experiment, a new set of participants (*n_older_* = 46, *n_younger_*= 50) transitioned directly from a baseline block to a counterbalanced, bidirectional 5.25° EC block.

Replicating observations from Experiment 2a, ECs evoked STA (1.93° ± 0.13°; main effect of EC Direction: *F(1, 94)* = 222.24, *p* = 1.68 x 10^-26^, *η^2^_p_* = 0.43, *Cohen’s F* = 1.54; *t(94)* = -14.91, *p* = 1.68 x 10^-26^, *d* = -3.07; **Fig. 3D**). The STA magnitude did not differ between age groups (main effect of Age Group: *F(1, 94)* = 0.39, *p* = 0.53; Age Group x Direction interaction: *F(1, 94)* = 0.10, *p* = 0.75). Notably, interpretation of these null results is more straightforward than for Experiment 2a, because overall movement angles were consistently directed towards the target (-0.41° ± 1.16°; main effect of Timepoint: *F(1, 92)* = 0.02, *p* = 0.90) in both age groups (main effect of Age Group: *F(1, 92)* = 0.29, *p* = 0.59; Age Group x Timepoint interaction: *F(1, 92)* = 0.40, *p* = 0.53; **Fig. 3E**).

Together, the results of Experiments 2a and 2b suggest that STA in response to small ECs with randomly varying directions does not differ between younger and older adults. Furthermore, any age-differences suggested by the numerical data involve a reduction of STA among older adults relative to younger adults, which would be an effect in the opposite direction of the age-related enhancement in adaptation reported here and elsewhere in the literature. This indicates that adaptation is not always enhanced among older adults, and suggests that exposure to bidirectional ECs reduces age-differences in adaptation. However, the results of Experiments 2a and 2b could also be explained if age-differences in adaptation are too small to observe via STA measurements.

### 3.3 Blocks of salient errors in opposite directions suppress age-differences in adaptation

To investigate whether variable errors undermine age-differences observed in adaptation that accumulates over a block of EC trials, we conducted Experiment 3. In this study, participants (*n_older_*= 50, *n_younger_* = 50) were exposed to two, 45° EC blocks. Both included errors in the same direction, but the second EC block followed a veridical feedback (FB) washout block that included salient errors in the opposite direction (**Fig. 4A-B**, **Supplementary Fig. 2A**). These opposite-direction errors resulted from the participant’s own adaptation in the preceding EC block (Avraham and Ivry, 2025). For example, consider a participant exposed to clockwise errors in the first EC block, leading to counterclockwise adaptation of movement angles. That movements are directed off-target would be revealed during the washout block with veridical FB, which shows the cursor at the position of the true mouse pointer, thus presenting the participant with visual errors in the direction opposite of the initial EC. This phenomenon allowed us to measure any changes in adaptation to a block of unidirectional ECs following exposure to errors in the opposite direction, while capturing age-differences in responses to the task.

**Figure 4.**
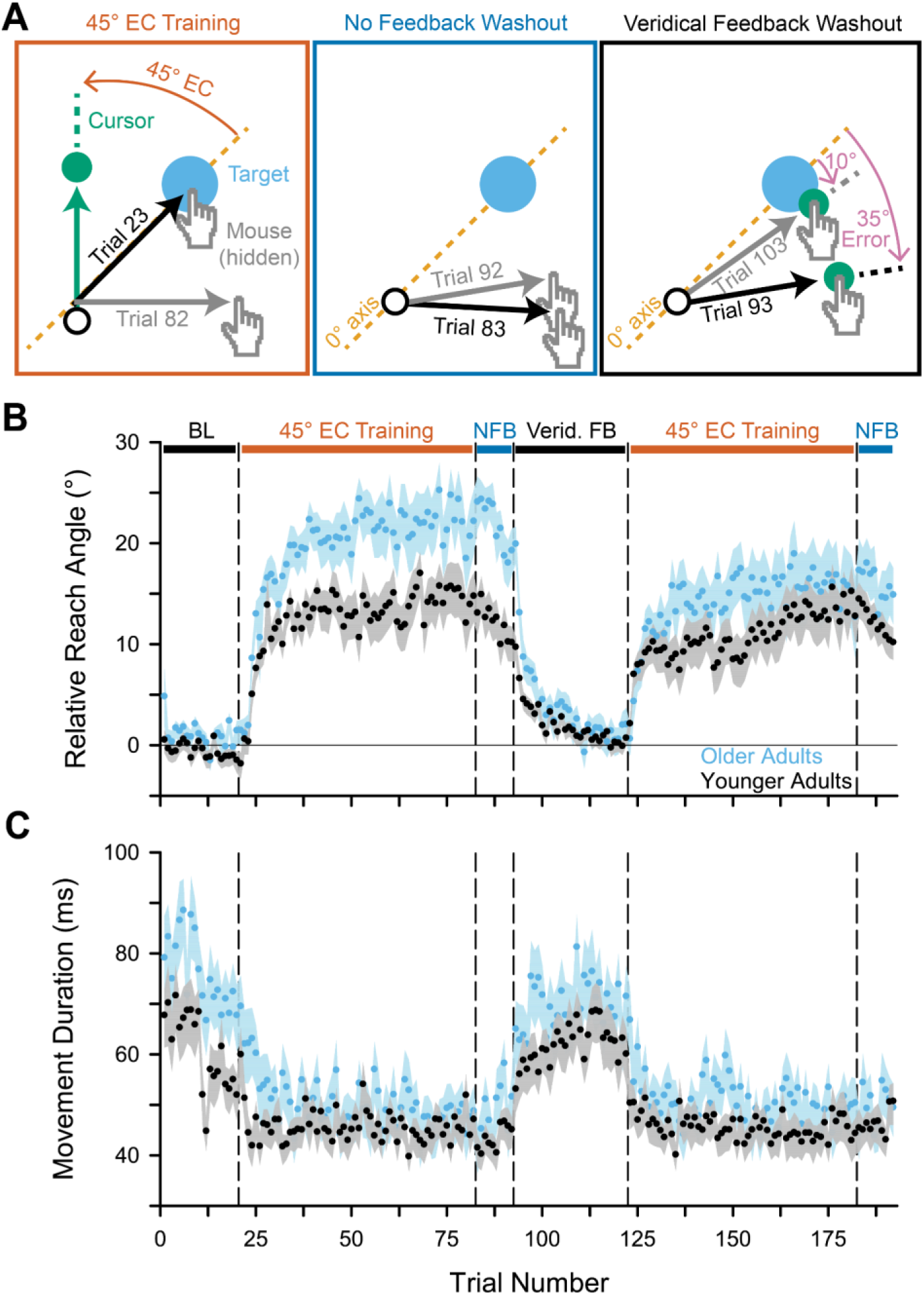
Implicit adaptation to unidirectional ECs before and after errors in the opposite direction. **A.** Schematic illustrating exemplar feedback provided to a during the EC, No-FB, and Washout phases of Experiment 3. **B.** Scatter plot showing the mean ± SEM movement angle on each trial of the experiment across participants. Movement direction is shown relative to the target and relative to the direction of the 45° EC, with positive relative movement direction values corresponding to the direction of movement opposite the direction of the error (*i.e.,* positive values indicate clockwise movements when the EC was in the counterclockwise direction). Data from older adults are shown in blue, and data from younger adults are shown in black. **C.** As in **B**, but showing movement durations. *Abbreviations*: EC: error-clamp, FB: feedback, SEM: standard error of the mean.

In line with our design, adapted movement angles persisted into the washout block (main effect of Timepoint: *F(4, 392)* = 101.24, *p_corrected_* = 3.34 x 10^-36^, *η^2^_p_* = 0.51, *Cohen’s F* = 1.02) for both younger (4.32° ± 0.52°, Baseline vs Washout: *t(98)* = 5.38, *p* = 2.74 x 10^-6^, *d* = 1.08) and older adults (7.66° ± 1.28°; *t(98)* = 7.35, *p* = 6.53 x 10^-10^, *d* = 1.48), ensuring that washout included errors opposite the direction of the ECs (**Fig. 4B**, **Supplementary Fig. 2A**). After the first 45° EC block, adaptation was greater among older adults (20.93° ± 2.12°) than younger adults (12.36° ± 1.68°; main effect of Age Group: *F(1, 98)* = 6.16, *p* = 0.01, *η^2^_p_* = 0.06, *Cohen’s F* = 0.25; *t(98)* = -3.17, *p* = 0.006, *d* = -0.64; **Fig. 4B**). Following the second 45° EC block, however, this age-difference dissipated (Age Group x Timepoint interaction: *F(4, 392)* =4.04, *p_corrected_* = 0.01, *η^2^_p_* = 0.04, *Cohen’s F* = 0.20; *t(98)* = -1.18, *p* = 0.33). Group movement angles converged because older adults exhibited less adaptation during the second 45° EC block (15.43° ± 2.43°; *t(98)* = -3.44, *p* = 0.003, *d* = -0.69) while adaptation remained stable among younger adults (12.12° ± 1.40°; *t(98)* = -0.151, *p* = 0.88).

Aging-related changes in the mechanisms of learning and memory may have caused this difference in responses to opposite-direction errors between age groups. However, older adults did exhibit greater adaptation during the first 45° EC block, leading to exposure to larger errors early on in the washout block (*t(98)* = -2.42, *p* = 0.04, *d* = -0.49) that may have more effectively suppressed adaptation in the second 45° EC block.

Nevertheless, the data clearly show that errors in the direction opposite of a subsequent EC suppresses adaptation among older adults, with the potential to reduce or eliminate age-differences in adaptation. Further supporting this conclusion, we did not observe any other differences in adaptation likely to provide alternate explanations to group differences in this experiment (**Supplementary Table 2**).

### 3.4 Age-differences in movement timing are unlikely to explain the effect of bidirectional errors on adaptation among older adults

We next analyzed whether the reduction in adaptation among older adults during the second 45° EC block of Experiment 3 could have resulted from changes in the time taken to move from the starting location past the target distance (movement duration) during the study. A change in movement duration would have changed the duration and timing of the EC, potentially affecting opportunities for the brain to adapt (Wang et al., 2024b). If older adults tended to move slower than younger adults in the first 45° EC block but caught up to younger adults’ movement speeds by the second 45° EC block, changes in movement duration could explain the results of Experiment 3 without appealing to differences in the adaptation process per se.

## 4. Discussion

Seemingly conflicting results in the literature indicated that implicit sensorimotor adaptation is not simply enhanced with age, and suggested that older adults experience different patterns of implicit learning depending on the kinds of errors generated in different tasks (Cisneros et al., 2026; Vandevoorde and Orban de Xivry, 2019; Hermans et al., 2025; Reuter et al., 2018). Here, we tested whether consistency in visual error, which differs between EC and force-field tasks, affected learning among older adults. Our data illustrate that the age-related enhancement of adaptation in an EC task depends on the persistence of a uniform error (Experiment 1, Section 3.1), and on the absence of errors in opposing directions (Experiments 2-3, Sections 3.2-3.3).

These findings uncover error variability as a proximal determinant of the degree of implicit motor adaptation among older adults. This addresses a puzzle in the literature, providing an explanation for discrepancies in results between different ways of querying implicit adaptation. Our findings further suggest that older adults’ implicit motor learning systems are resilient to aging-related changes under day-to-day conditions with normally distributed errors, highlighting a natural mechanism for healthy aging of the implicit motor system.

### 4.1. Possible cognitive and neural mechanisms

Based on previous literature, we favor two kinds of cognitive and neural mechanisms for age differences in the patterns of adaptation expressed in response to variable errors. Error variability could suppress implicit adaptation among older adults through either mechanism, or through a combination of both.

The first possible mechanism involves the acquisition of conflicting memories of implicit adaptation that suppress the net behavioral expression of that adaptation (destructive interference). The second mechanism appeals to the possibility that participants’ meta-knowledge of the task could affect sensory processing of error signals and affect downstream learning.

#### 4.1.1. Destructive Interference

Training with variable errors could have reduced the extent of observable adaptation among older adults by generating conflicting implicit memories. Implicit adaptation to errors in one direction interferes destructively with learning in response to errors in the opposite direction among younger adults. This interference leads to net-zero learning in paradigms with errors counterbalanced throughout (*e.g.*, Howard et al., 2013; Sheahan et al., 2016; Kim et al., 2022), as in Experiment 2. In studies with block designs like Experiment 3, destructive interference reduces the magnitude of learned behavioral change upon re-exposure to an error (anti-savings; Avraham and Ivry, 2025). Such anti-savings is theorized to arise from simultaneously activated cerebellar memories that drive movements in opposite directions (Hanajima et al., 2015; Raymond and Medina, 2018; Shlerf et al., 2012; Wang and Ivry, 2025).

Cerebellar memory recall is influenced by error history (Heald et al., 2021, but see Wang et al., 2024a) and by pattern separation computations in granule neurons (Cayco-Gajic and Silver, 2019; Nguyen et al., 2023). These neurons degenerate as we age (Andersen et al., 2003; Arleo et al., 2024), potentially leading to reduced pattern separation and greater generalization of cerebellar learning among older adults. Therefore, age-related reductions in granule cell pattern separation could result in greater recall of cerebellar memories established from different directions of error, reducing behavioral expression of motor learning through destructive memory interference.

In the context of our study, impaired pattern separation could have led to greater interference between clockwise and counterclockwise adaptation in response to different error directions among older adults. This would explain why we observed similar levels of adaptation between groups after the second block of EC feedback in Experiment 3 despite enhanced initial adaptation among older adults (**Fig. 4**). In this case, future neuroimaging studies could reveal key evidence about differences in cerebellar activity that underlie age-related changes in implicit adaptation and anti-savings, even at times when behavioral expression of adaptation is similar between older and younger adults.

#### 4.1.2. Metacognitive Effects on Sensory Processing

A second possible mechanism explaining our results involves interactions between implicit adaptation and participants’ explicit knowledge about their movements. Participants exposed to consistent ECs may have lacked insight into the variability of their own movements, as the task instructions involve an explanation about the nature of the perturbation but not the nature of typical movements. However, observation of the the cursor moving in the opposite direction of previous ECs during a veridical FB phase (Experiment 3) or a block of mixed direction ECs (Experiment 2), could have led participants to recognize that the degree of stability in the EC feedback does not necessarily reflect stability in their movements.

After recognizing this additional layer of deception inherent to the EC, participants may have attempted to increase their movement accuracy on EC trials by directing their attention away from visual information and towards proprioceptive information. Because differences in visuo-proprioceptive integration during blocks of consistent ECs may contribute to age-related enhancement of implicit adaptation (Cisneros et al., 2026; Kitchen and Miall, 2021; Van De Plas and Orban de Xivry, 2026; Carranza et al., 2025), awareness-driven changes in the weights assigned to visual and proprioceptive information may have mitigated age-related differences in learning. Indeed, attentional shifts away from visual FB limit the amount of adaptation that visual errors, like ECs, can evoke while changing visuo-proprioceptive integration at the neural level (Block and Bastian, 2010; Hsiao et al., 2022). In other words, variability in visual FB may increase reliance on proprioceptive information, and this change may be particularly impactful among older adults who had previously downweighted proprioceptive information.

### 4.2. Caveats and Limitations

While our study highlights error variability as a key factor influencing implicit adaptation in older adults through multiple possible mechanisms, some key limitations caution against overinterpretation of our data. Firstly, our reliance on highly artificial EC manipulations leaves open questions about how older adults learn in response to more naturalistic stimuli. Secondly, discrepancies between our data and a previous report leave the effect of variable visual error presentation on implicit adaptation among young adults unclear. Third, our experimental design leaves open questions about what kinds of error variability influence implicit adaptation among older adults.

#### 4.2.1. Artificiality of the EC Manipulation

In the current study, we imposed errors via ECs, which are unlikely to emulate natural errors. So, whether our findings generalize to situations encountered in everyday life remains an open question. Nevertheless, the EC paradigm offered a key advantage: complete control over the visual errors participants experienced. This allowed us to test how adaptation differed in response to identical visual feedback between older and younger adults, and between older adults with different experiences with directional errors. Because participants experienced identical errors during EC blocks, we can infer that any differences in learning between age groups are attributable to differences in the processing of those perturbations, as opposed to differences in the specific errors that the perturbation evoked from participants with variable behavioral responses.

#### 4.2.2. Effects of error variability on adaptation in young adults

Based on our control of the errors that participants experienced, we are confident that all group differences observed in Experiments 1 and 2 are attributable to the groups’ age differences. However, we observed a group difference in anti-savings following opposite-direction errors in Experiment 3 that we cannot clearly attribute to age. In that experiment, opposite-direction errors were imposed through trials with veridical feedback, rather than ECs. This led to opposite-direction errors of greater magnitude and variability for older adults, who exhibited anti-savings, as compared to younger adults, whose adaptation was not statistically significantly affected by exposure to opposite-direction errors (**Supplementary Fig. 2**, **Fig. 4**). Therefore, it is possible that younger adults may have exhibited anti-savings if provided with the same errors as the older adults in our study. Indeed, the report that Experiment 3 was inspired by (Avraham & Ivry, 2025) evoked anti-savings among younger adults trained for about 4x as long as those in our task, indicating that younger adults can exhibit anti-savings with more intensive training conditions. Regardless of difficulties in interpreting our lack of an effect among younger adults, we can nevertheless conclude that exposure to opposite-direction errors blunted adaptation among older adults in our study.

#### 4.2.3. Influence of Variability in Error Direction Versus Magnitude

Although our experiments generally emphasized manipulations of error direction, we cannot rule out the possibility that adaptation among older adults depends on variability in error magnitude. Changes in error magnitude and direction were not temporally separated in Experiment 3. Experiment 2’s results suggest that variability in error direction is sufficient to prevent age-related amplification of implicit adaptation, but they do not rule out a role for variability in error magnitude. Additional follow-up studies will be required to evaluate whether consistency in the error direction is key, or whether consistency in error magnitude is also necessary to amplify adaptation among older adults.

### 4.3. Broader Implications

Regardless of whether the key mechanism lies in variability of error magnitude or direction, our data clearly show that error variability limits implicit adaptation among older adults. Because one would generally expect errors to vary in day-to-day life, with errors in movements falling in something like a normal distribution around zero for healthy adults, our results suggest that older and younger adults are likely to exhibit similar levels of adaptation under typical circumstances. In other words, older adults have a natural system that can counteract changes in adaptation that emerge with age, protecting them from hyperrecruitment of implicit systems that change behavior without their conscious awareness. Nevertheless, that enhanced implicit adaptation among older adults requires a consistent perturbation suggests that practitioners could leverage carefully planned physical therapy interventions (*e.g.*, in virtual reality) that are not subject to the statistics of the natural environment to increase the gain of implicit contributions to movement among patients.

We speculate that patterns of implicit adaptation in response to ECs may be similar to adaptation driven by other forms of errors that drive cerebellar learning. For example, errors driving visuomotor adaptation are encoded by olivo-cerebellar pathways parallel to those driving learning in response to pain (Ju et al., 2019; Titley et al., 2020; Miles and Wiesendanger, 1975; Kitazawa et al., 1998; Lisberger, 2021; Wang and Ivry, 2025). This suggests that injuries resulting in consistent pain could evoke stronger implicit motor adaptation among older adults relative to younger adults, putting older people at greater risk of subconsciously developing postural adaptations to pain that cause them other injury-related complications down the line. Future work will be necessary to determine whether consistent pain-related errors could cause implicit adaptation that affects hand/arm movement differently among younger and older adults.

## Declaration of generative AI and AI-assisted technologies in the manuscript preparation process

We used Google Gemini to refine and debug python scripts for data analysis and to improve the organization of the methods section. The authors reviewed all suggestions from Gemini for accuracy and suitability for this project before accepting or rejecting them.

## Supplementary Figures

**Supplementary Figure 1.**
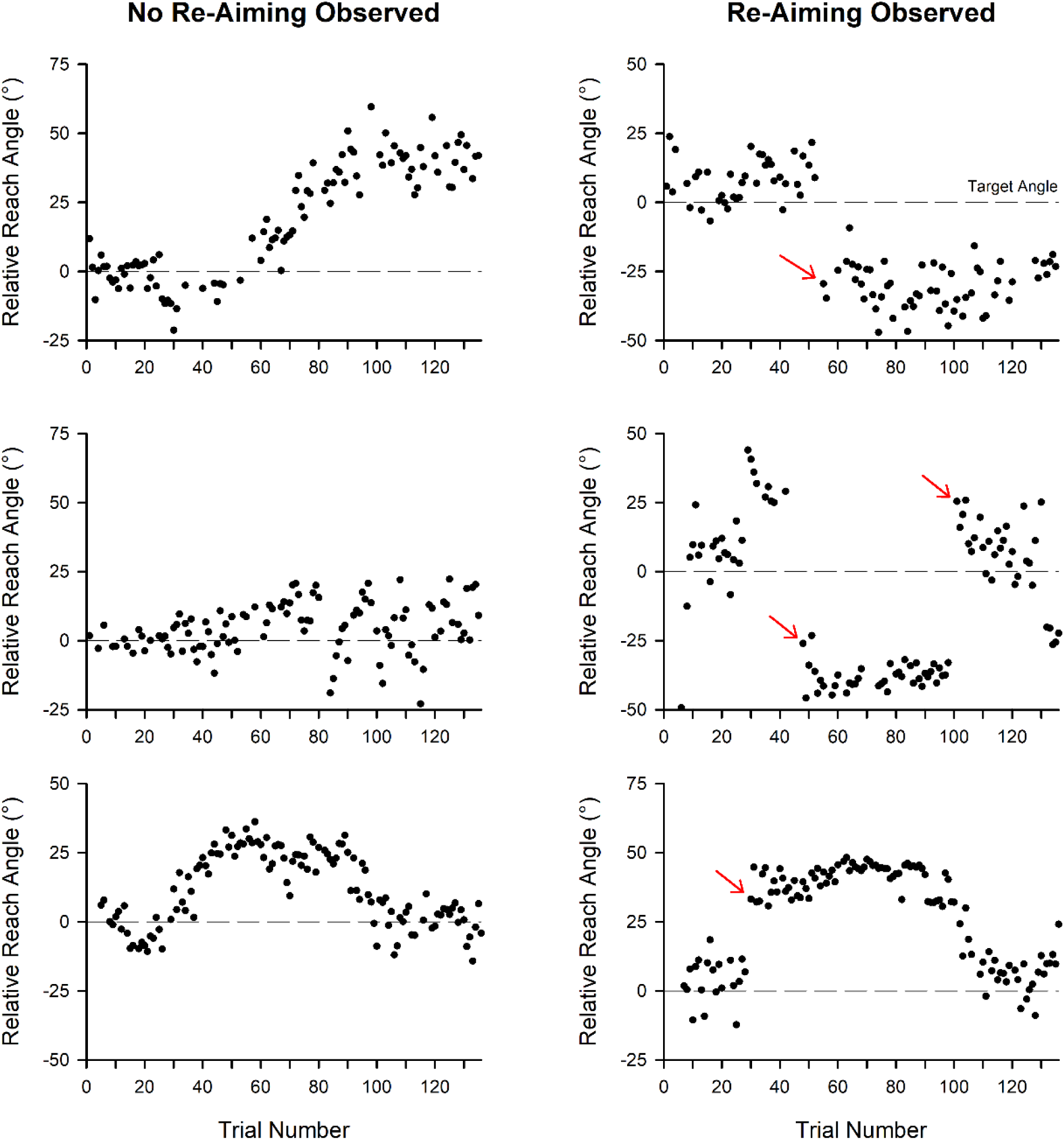
Movement directions for example participants. Scatter plots showing movement direction on each trial during Experiments 1 and 3. Data from participants included in the final analysis are shown at left, and data from participants with automatically-detected re-aiming are shown at right. Red arrows indicate the trial on which a sustained re-aiming bout was flagged.

**Supplementary Figure 2.**
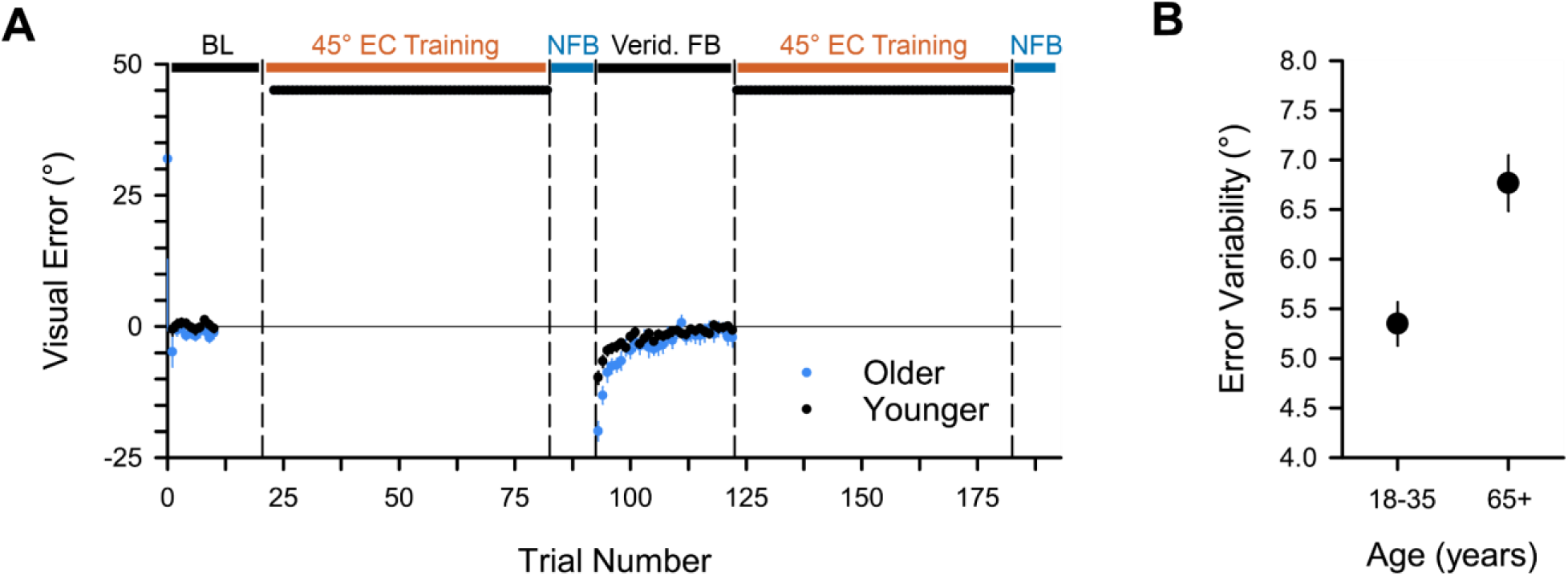
Errors shown during Experiment 3. **A.** Scatter plot showing the mean ± SEM error angle on each trial of the experiment across participants. Error angle is shown relative to the target and relative to the direction of the 45° EC, with positive relative error angle values corresponding to the direction of the error during the EC block (*i.e.,* negative values indicate counterclockwise movements when the EC was clockwise). Data from older adults are shown in black, and data from younger adults are shown in red. **B.** As in **A**, but showing the mean ± SEM error variability for each group during the washout phase of the study. There was a significant difference in error variability between groups (*t*(98) = 3.95, *p* = 0.0001). *Abbreviations*: EC: error-clamp, FB: feedback, SEM: standard error of the mean.

## Supplementary Tables

**Supplementary Table 1.** Participant handedness.

| Exp. | Group | Right Handed,<br>Used Right | Right Handed,<br>Used Left | Left Handed,<br>Used Right | Left Handed,<br>Used Left | Ambidextrous,<br>Used Right | Ambidextrous,<br>Used Left |
|---|---|---|---|---|---|---|---|
| 1 | Younger | 41 | 0 | 4 | 2 | 0 | 0 |
|  | Older | 21 | 0 | 2 | 1 | 1 | 0 |
| 2a | Younger | 41 | 0 | 4 | 2 | 0 | 0 |
|  | Older | 20 | 0 | 2 | 0 | 1 | 0 |
| 2b | Younger | 46 | 0 | 3 | 1 | 0 | 0 |
|  | Older | 41 | 0 | 1 | 1 | 3 | 0 |
| 3 | Younger | 49 | 0 | 1 | 0 | 0 | 0 |
|  | Older | 46 | 0 | 1 | 2 | 1 | 0 |

**Supplementary Table 2.** Pre-planned post-hoc pairwise comparisons of movement angle across timepoint and age-group in Experiment 3.

| Comparison | t | df | p<sub>adj</sub> | d |
|---|---|---|---|---|
| Older vs Younger: Baseline | -1.92 | 98 | 0.10 | -0.39 |
| Older vs Younger: First No FB Block | -3.12 | 98 | **0.006** | -0.64 |
| Older vs Younger: Early Washout | -2.42 | 98 | **0.04** | -0.49 |
| Older vs Younger: Late Washout | -0.86 | 98 | 0.43 | -0.17 |
| Older vs Younger: Second No FB Block | -1.18 | 98 | 0.33 | -0.24 |
| Older: Baseline vs Early Washout | 7.35 | 98 | **6.53 x 10<sup>-10</sup>** | 1.48 |
| Older: Baseline vs Late Washout | 1.11 | 98 | 0.33 | 0.22 |
| Older: First vs Second No FB Block | -3.44 | 98 | **0.003** | -0.69 |
| Younger: Baseline vs Early Washout | 5.38 | 98 | **2.74 x 10<sup>-6</sup>** | 1.09 |
| Younger: Baseline vs Late Washout | 1.61 | 98 | 0.17 | 0.32 |
| Younger: First vs Second No FB Block | -0.15 | 98 | 0.88 | -0.03 |

*Note.* The post-hoc tests for the mixed-design ANOVA model used here involve the same number of degrees of freedom (df) for between-subjects and within-subjects comparisons because the underlying ANOVA was constructed as a multivariate linear regression. This involves pooling the covariance matrix and globally estimating degrees of freedom from the shared covariance matrix. This approach does not rely on the assumption of sphericity (Maxwell et al., 2018; Singmann et al., 2025), which is violated by our data, making it more robust than post-hoc tests conducted on the raw data. P-values were adjusted according to the Benjamini-Hochberg procedure for controlling the False Discovery Rate.

**Supplementary Table 3.** Post-hoc pairwise comparisons of movement duration across timepoints in Experiment 3.

| Timepoint | Age-Group | contrast | df | t | p<sub>adj</sub> | d |
|---|---|---|---|---|---|---|
| Baseline | . | older - younger | 98 | 3.22 | **0.004** | 0.65 |
| First 45° EC | . | older - younger | 98 | 1.25 | 0.30 | 0.25 |
| First no-FB | . | older - younger | 98 | 1.37 | 0.27 | 0.28 |
| Washout | . | older - younger | 98 | 1.74 | 0.16 | 0.35 |
| Second 45° EC | . | older - younger | 98 | 1.39 | 0.27 | 0.28 |
| Second no-FB | . | older - younger | 98 | 0.97 | 0.42 | 0.20 |
| . | older | Baseline - First 45° EC | 98 | 7.55 | **4.18 x 10<sup>-10</sup>** | 1.52 |
| . | older | Baseline - First no-FB | 98 | 7.57 | **4.18 x 10<sup>-10</sup>** | 1.53 |
| . | older | Baseline - Washout | 98 | 0.49 | 0.70 | 0.10 |
| . | older | Baseline - Second 45° EC | 98 | 6.55 | **1.62 x 10<sup>-8</sup>** | 1.32 |
| . | older | Baseline - Second no-FB | 98 | 5.87 | **2.64 x 10<sup>-7</sup>** | 1.19 |
| . | older | First 45° EC - First no-FB | 98 | 0.96 | 0.42 | 0.19 |
| . | older | First 45° EC - Washout | 98 | -6.94 | **3.01 x 10<sup>-9</sup>** | -1.40 |
| . | older | First 45° EC - Second 45° EC | 98 | -0.39 | 0.76 | -0.08 |
| . | older | First 45° EC - Second no-FB | 98 | -0.26 | 0.84 | -0.05 |
| . | older | First no-FB - Washout | 98 | -7.21 | **1.04 x 10<sup>-9</sup>** | -1.46 |
| . | older | First no-FB - Second 45° EC | 98 | -1.29 | 0.29 | -0.26 |
| . | older | First no-FB - Second no-FB | 98 | -0.96 | 0.42 | -0.19 |
| . | older | Washout - Second 45° EC | 98 | 7.26 | **1.04 x 10<sup>-9</sup>** | 1.47 |
| . | older | Washout - Second no-FB | 98 | 5.96 | **2.07 x 10<sup>-7</sup>** | 1.20 |
| . | older | Second 45° EC - Second no-FB | 98 | 0.006 | 0.99 | 0.001 |
| . | younger | Baseline - First 45° EC | 98 | 3.03 | **0.007** | 0.61 |
| . | younger | Baseline - First no-FB | 98 | 3.56 | **0.002** | 0.72 |
| . | younger | Baseline - Washout | 98 | -2.06 | 0.08 | -0.42 |
| . | younger | Baseline - Second 45° EC | 98 | 2.73 | **0.02** | 0.55 |
| . | younger | Baseline - Second no-FB | 98 | 2.10 | 0.08 | 0.42 |
| . | younger | First 45° EC - First no-FB | 98 | 1.39 | 0.27 | 0.28 |
| . | younger | First 45° EC - Washout | 98 | -5.29 | **2.42 x 10<sup>-6</sup>** | -1.07 |
| . | younger | First 45° EC - Second 45° EC | 98 | 0.06 | 0.98 | 0.01 |
| . | younger | First 45° EC - Second no-FB | 98 | -0.54 | 0.68 | -0.11 |
| . | younger | First no-FB - Washout | 98 | -5.85 | **2.64 x 10<sup>-7</sup>** | -1.18 |
| . | younger | First no-FB - Second 45° EC | 98 | -1.31 | 0.29 | -0.26 |
| . | younger | First no-FB - Second no-FB | 98 | -1.55 | 0.22 | -0.31 |
| . | younger | Washout - Second 45° EC | 98 | 5.73 | **4.00 x 10<sup>-7</sup>** | 1.16 |
| . | younger | Washout - Second no-FB | 98 | 4.32 | **0.0001** | 0.87 |
| . | younger | Second 45° EC - Second no-FB | 98 | -0.81 | 0.51 | -0.16 |

*Note*: The post-hoc tests for the mixed-design ANOVA model used here involve the same number of degrees of freedom (df) for between-subjects and within-subjects comparisons because the underlying ANOVA was constructed as a multivariate linear regression. This involves pooling the covariance matrix and globally estimating degrees of freedom from the shared covariance matrix. This approach does not rely on the assumption of sphericity (Maxwell et al., 2018; Singmann et al., 2025), which is violated by our data, making it more robust than post-hoc tests conducted on the raw data. P-values were adjusted according to the Benjamini-Hochberg procedure for controlling the False Discovery Rate.

## Notes

### Competing Interest Statement

The authors have declared no competing interest.

